# A HaloTag–4R-Tau Pulse-Chase Sensor Reveals Neddylation Inhibition Promotes Degradation of Tau in iNeurons

**DOI:** 10.64898/2026.01.28.702105

**Authors:** Qiang Xiao, Zi Gao, Seth Allen, Danny Garza, Richard I. Morimoto, Jeffery W. Kelly

## Abstract

Tau accumulation is a central driver of neurodegenerative diseases, yet strategies to promote its clearance remain limited. We developed a HaloTag–4R-Tau sensor in human iPSC-derived neurons (iNeurons) that enables sensitive monitoring the kinetics of both lysosomal partitioning and overall cellular turnover of tau. Using this sensor, we screened a small collection of small-molecule modulators of proteostasis network function and identified Neddylation inhibition by Pevonedistat as a robust promoter of soluble tau degradation. Mechanistic analysis including proteomic profiling revealed that Neddylation inhibition hastens HaloTag-Tau clearance via compensatory activation of a proteasome-dependent pathway(s) as well as the autophagy-lysosome pathway. Our findings establish a powerful tool for probing tau homeostasis and highlight Neddylation inhibition as a potential therapeutic approach for enhancing both proteasome and lysosome-mediated tau clearance in tauopathies.

## Introduction

Abnormal accumulation of the tau protein in neurons, by way of misassembly into neurofibrillary tangles (NFTs) and additional aggregate structures, is a pathological hallmark shared by Alzheimer’s Disease (AD), Frontotemporal Dementia (FTD), and other neurodegenerative disorders^1,2^. Multiple therapeutic strategies—including antibodies against tau NFTs and pharmacological inhibition of tau post-translational modifications—have been evaluated recently in clinical trials; however, none of these drug candidates have been approved for the treatment of tau pathology due to lack of efficacy or on-target side effects^3^. More recently, advances in antisense oligonucleotide (ASO) therapies have shown promise in clearing tau NFTs from the brains of AD patients by reducing tau translation^4^. A clinical trial highlighted the feasibility of clearing tau NFTs by lowering soluble tau levels through intrathecal (IT) administration of an ASO^4^. Identifying targets that facilitate soluble tau clearance is an alternative therapeutic strategy for tauopathies. To date, such efforts have not yielded any clinical candidates, largely due to the lack of sufficiently sensitive tools^5^ or because investigations were performed primarily in non-human neuronal models^6^.

To address this gap, we developed a pulse chase-based HaloTag–4R-Tau sensor in established NGN2-induced iPSC-derived neurons (iNeurons)^7^ to identify targets that, when pharmacologically modulated, hasten tau clearance. We previously established that a commonly used protein tag, HaloTag, becomes proteolytically resistant within lysosomes after it reacts with its small molecule stabilizer ligand fused to a Tetramethylrhodamine (TMR) fluorophore. Hastened time-dependent buildup of TMR-HaloTag enables activation of autophagy to be quantified by a pulse-chase experiment^8^. About 95% of TMR–HaloTag is degraded by the proteasome, but this is reduced to ≈ 90% upon autophagy activation—thus both lysosomal and proteasomal degradation can be monitored by this tag by quantification of its band intensity via SDS-PAGE in-gel fluorescence of TMR^8^. In the present study, we leveraged this feature by fusing HaloTag to 4R-tau, allowing us to track HaloTag–4R-Tau translocation from the soma to lysosomes as an indicator of lysosomal tau degradation, as well as to measure the overall cellular tau turnover by quantifying the in-gel fluorescence intensity of the TMR-HaloTag-4RTau by SDS-PAGE. Tau fused to the C-terminus of HaloTag has a normal subcellular localization and an analogous half-life to endogenous Tau in neurons.

Using TMR-HaloTag-4RTau, we screened a library of pharmacological modulators of the proteostasis network, given that disruption of proteostasis network function is a major driver of pathological protein aggregation in various neurodegenerative diseases, including tau NFT formation^9^. We identified compounds that either promoted or inhibited lysosomal localization and/or cellular turnover of TMR-HaloTag-4RTau. The pathways modulated by this pharmacology provides insight into how tau homeostatsis is regulated by the proteostasis network. Neddylation inhibition emerged as a strategy for hastening soluble tau degradation. Mass spectrometry proteomic analysis demonstrated that Neddylation inhibition-mediated tau clearance likely involves both the autophagy-lysosome pathway and, predominantly, the ubiquitin-proteasome system.

## Results

### HaloTag-Tau sensor allows detection of its lysosomal translocalization in iNeurons

We used CRISPR to knock in NGN2, a transcription factor that drives neuronal differentiation from stems cells^10^, into the CLYBL safe-harbor locus of a commercially available iPSC line, following a published protocol^7^ (Figure 1A). After subcloning, the NGN2-engineered iPSCs (with normal karyotyping, Figure S1A) were infected with lentivirus encoding HaloTag or HaloTag–4R-Tau. Following antibiotic selection, surviving iPSCs were differentiated into glutamatergic excitatory cortical neurons by turning on expression of NGN2 by adding Doxycycline (Dox). After a two-week differentiation, we confirmed the expression of key neuronal makers, including NeuN, MAP2, and TUJ-1, by immunostaining (Figure S1B). Once the neurons were fully mature, HaloTag or HaloTag– 4RTau were labeled with the HaloTag ligand, TMR, allowing visualization of HaloTag and HaloTag-4RTau within the cortical neurons employing live-cell imaging (Figure 1B). As shown in Figure 1C (first column), HaloTag was ubiquitously distributed within the soma, whereas HaloTag-4R-TauP301L was preferentially localized to somatic cytoskeleton and axons, indicating the fusion of HaloTag does not interrupt tau’s microtubule binding properties in cortical neurons. We also validated that this microtubule binding property was not interrupted by the fusion to HaloTag, as evidenced by its colocalization with immunostaining of microtubules in neuroblastoma cells (Figure S1C).

**Figure 1.**
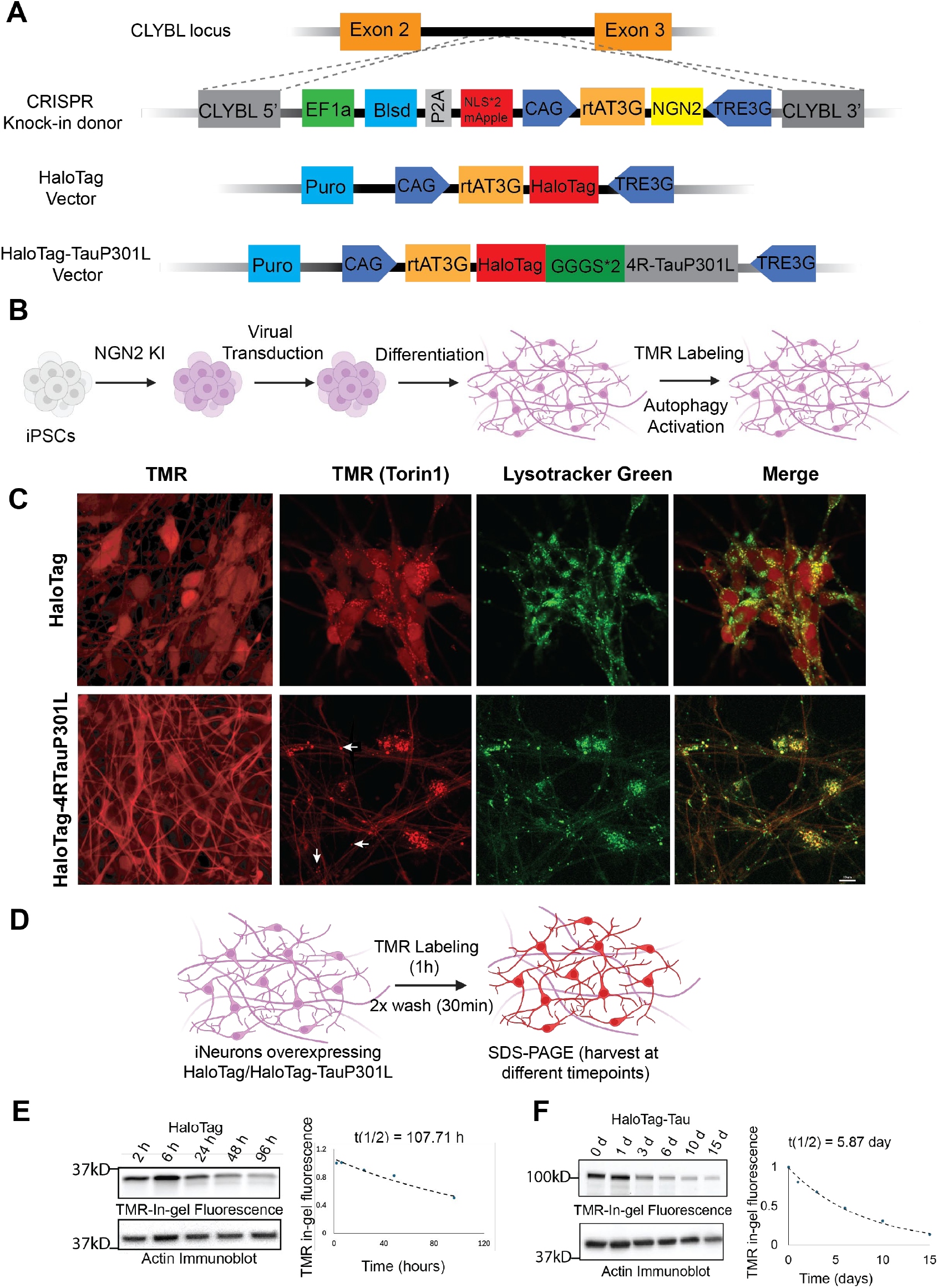
HaloTag–Tau sensor enables visualization of tau translocation and measurement of its half-life in iNeurons. (A) Schematic of constructs used: NGN2 plasmid for CRISPR knock-in at the CLYBL locus, and inducible lentiviral vectors encoding HaloTag or HaloTag–Tau. (B) Workflow for generating iNeurons with the HaloTag/HaloTag–Tau sensor and performing drug treatments. iPSCs were subcloned following NGN2 knock-in and enriched by puromycin selection after lentiviral transduction. HaloTag/HaloTag–Tau was labeled with TMR, followed by 24-hour treatment with Torin1 (500 nM) to activate autophagy. (C) Representative live-cell images of iNeurons expressing HaloTag or HaloTag–Tau after TMR labeling. Cells were treated with DMSO (TMR column) or Torin1 (500 nM, TMR (Torin1) column) for 24 hours. Lysotracker Green (50 nM) was added after Torin1 treatment (Lysotracker Green column), and merged images show colocalization of TMR and Lysotracker channels. Scale bar, 10 μm. (D) Schematic of iNeuron culture and labeling paradigm for half-life measurement of HaloTag and HaloTag– Tau. (E–F) SDS–PAGE analysis of lysates from iNeurons expressing HaloTag or HaloTag–Tau harvested at different time points, with quantification plots (n = 3). Half-lives were calculated by fitting the data to an exponential decay function.

We next treated the iNeurons with an mTOR inhibitor, Torin 1 (500nM), a well-established autophagy activator. After a 24 hour-treatment, we observed a marked accumulation of puncta derived from both HaloTag and HaloTag–4RTau within neuronal soma (predominantly) and axons (as indicated with white arrows, Figure 1C Column 2). These puncta showed strong colocalization with lysotracker, a lysosomotrophic dye, indicating lysosomal localization (Figure 1C, columns 3-4). To further assess whether lysosomal localization of HaloTag–4RTau is autophagy dependent, we examined the effector function of key autophagy genes, including ATG5 and SQSTM1 (p62). We transduced the same HaloTag-4RTau construct (Figure 1A) into U251 cells (wild-type and ATG5 knockout) and HEK293 cells (wild-type and p62 knockout), followed by autophagy activation by Torin1. As shown in Figure S1D-E and Figure S1F-G, ATG5 knockout completely abolished lysosomal localization of HaloTag–Tau, while p62 knockout significantly reduced its lysosomal accumulation. These results demonstrate that lysosomal targeting of HaloTag–4RTau is dependent on the autophagy pathway.

We next performed a pulse–chase experiment (Figure 1D) to determine the half-life of HaloTag or HaloTag–4R-Tau using SDS–PAGE. Briefly, HaloTag or HaloTag–4R-Tau in iNeurons was labeled with HaloTag ligand TMR for 1 hour, followed by 2 media washes over the period of 30 minutes, and the fluorescence abundance of HaloTag or HaloTag–4R-Tau band was monitored at multiple time points by SDS–PAGE analysis in cell lysates collected over time. The half-life of HaloTag was approximately 4.5 days (Figure 1E), whereas HaloTag–Tau exhibited a half-life of approximately 6 days (Figure 1F). These results indicate that the HaloTag–Tau sensor undergoes degradation kinetics similar to endogenous tau, whose half-life has been reported to be ∼6.7 days^11^.

### Neddylation inhibition promotes lysosomal localization and cellular turnover of HaloTag-Tau

Using lysosomal localization of the HaloTag–4RTau sensor in iNeurons as a phenotypic readout (Figure 2A), we screened a small-molecule library targeting the function of diverse pathways within the proteostasis network (https://www.proteostasisconsortium.com/proteostasis-regulator-plates/). We identified compounds that either accelerated or inhibited lysosomal localization of HaloTag–4RTau after 24 h of pharmacological treatment, as summarized in Figure 2B (see Figure S2A for the heatmap with hit annotations). In addition to canonical autophagy inhibitors, inhibition of HSP90, inositol-requiring enzyme 1 (IRE1), USP10, PIKfyve, and calcium channels almost completely abolished lysosomal localization of HaloTag–4RTau (Figure S2B). Among these hits, inhibition of USP10 has been reported to suppress autophagy^12^, whereas the effects of HSP90, IRE1 and PIKfyve inhibition on autophagy are context-dependent^13-16^. These observations suggest that modulation of proteostasis network function can influence lysosomal degradation of HaloTag-4RTau either directly or indirectly. In addition to mTOR inhibitors, we identified A-485, a histone acetyltransferase inhibitor, and Pevonedistat, a neddylation inhibitor, as strong inducers of HaloTag-Tau lysosomal localization (Figure 2C-D), consistent with previous reports that inhibition of histone acetyltransferase or neddylation activates autophagy^17, 18^.

**Figure 2.**
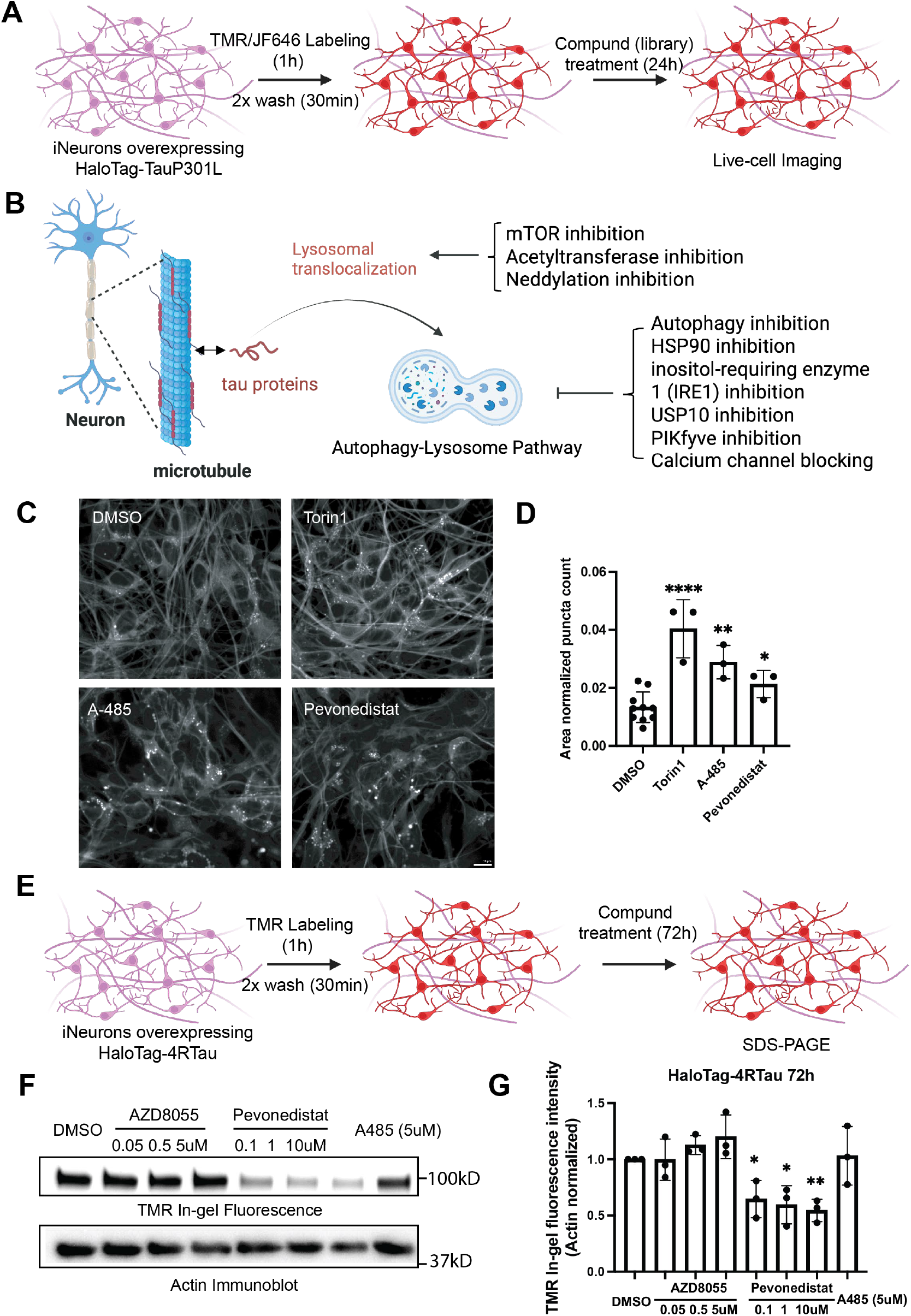
HaloTag–Tau sensor enables high-content screening of proteostasis modulators to assess effects on lysosomal translocation of Tau. (A) Schematic of the experimental workflow for treating iNeurons expressing the HaloTag–Tau sensor with a library of proteostasis modulators for live-cell imaging. (B) Summary of pathways of hits that either promote or inhibit lysosomal translocation of Tau. (C) Representative live-cell images of vehicle control and compounds that promote lysosomal localization of HaloTag–Tau. All drugs were applied at 10 μM in triplicate wells. Scale bar, 5 μm. (D) Quantification of positive hits based on the number of puncta normalized to neuronal area. Each drug was tested in three independent wells (n = 3). Data are presented as mean ± SD. Statistical significance was determined using an unpaired Student’s t-test: *p < 0.05, **p < 0.01, ****p < 0.0001. (E) Schematic of the experimental workflow for treating iNeurons expressing the HaloTag–Tau sensor with AZD8055, Pevonedistat, and A-485. (F) Representative full SDS–PAGE in-gel fluorescence images and immunoblots of TMR–HaloTag–Tau under different treatment conditions. (G) Quantification and statistical analysis of band intensities of full-length TMR–HaloTag–Tau from in-gel fluorescence. Data are presented as mean ± SD. Statistical significance was determined using an unpaired Student’s t-test: *p < 0.05, **p < 0.01.

We then examined Tau clearance capacity of those identified hits by running the same pulse-chase experiment in iNeurons overexpressing HaloTag-4RTau (Figure 2E) with treatment of an mTOR inhibitor, AZD8055, Neddylation inhibitor, Pevonedistat, or Histone acetyltransferase inhibitor, A-485, for 72 hours. The cell lysates were then harvested for SDS-PAGE analysis. In-gel TMR-HaloTag-4RTau fluorescence intensity (normalized to housekeeping protein Actin) was quantified as indicative of HaloTag-4RTau abundance. As shown in Figure 2F-G, all three doses of Pevonedistat significantly decrease the full-length HaloTag-4RTau band intensity, whereas mTOR inhibitor and A-485 failed to do so at tested concentrations. These results demonstrate that Neddylation inhibition mediated HaloTag-4RTau clearance is more efficient than mTOR-inhibition or A-485 mediated autophagy-lysosome pathway activation, suggesting that the means by which Pevonedistat activates autophagy is well suited to HaloTag-4RTau degradation and that Neddylation inhibition mediated HaloTag-4RTau clearance may occur by lysosomal and proteosomal degradation.

### Both the autophagy-lysosome pathway and ubiquitin-proteasome system are responsible for HaloTag-4RTau turnover

To investigate whether the proteasome or autophagy–lysosome pathway is primarily responsible for the clearance of HaloTag–Tau under basal conditions, we performed a pulse–chase experiment using the HaloTag–4R-Tau sensor in neuroblastoma cells (Figure 3A). Cells were treated with an autophagy inhibitor, including a ULK1 inhibitor (MRT68921), VPS34 inhibitor (SAR405), v-ATPase inhibitor (Bafilomycin A1), or a proteasome inhibitor (marizomib), or the combination of an autophagy inhibitor and proteasome inhibitor for 24 hours. As shown in Figure 3B-C, autophagy inhibitors MRT68921 and Bafilomycin A1 slightly increased the band intensity of full-length HaloTag-4RTau, whereas the proteasome inhibitor substantially enhanced the band intensity. Surprisingly, the combination of proteasome inhibitor with Bafilomycin A1 led to a significant further enhancement of the full-length HaloTag-4RTau band intensity, suggesting both proteasome and autophagy-lysosome pathways are responsible for HaloTag-Tau turnover, consistent with prior Tau degradation conclusions^19^.

**Figure 3.**
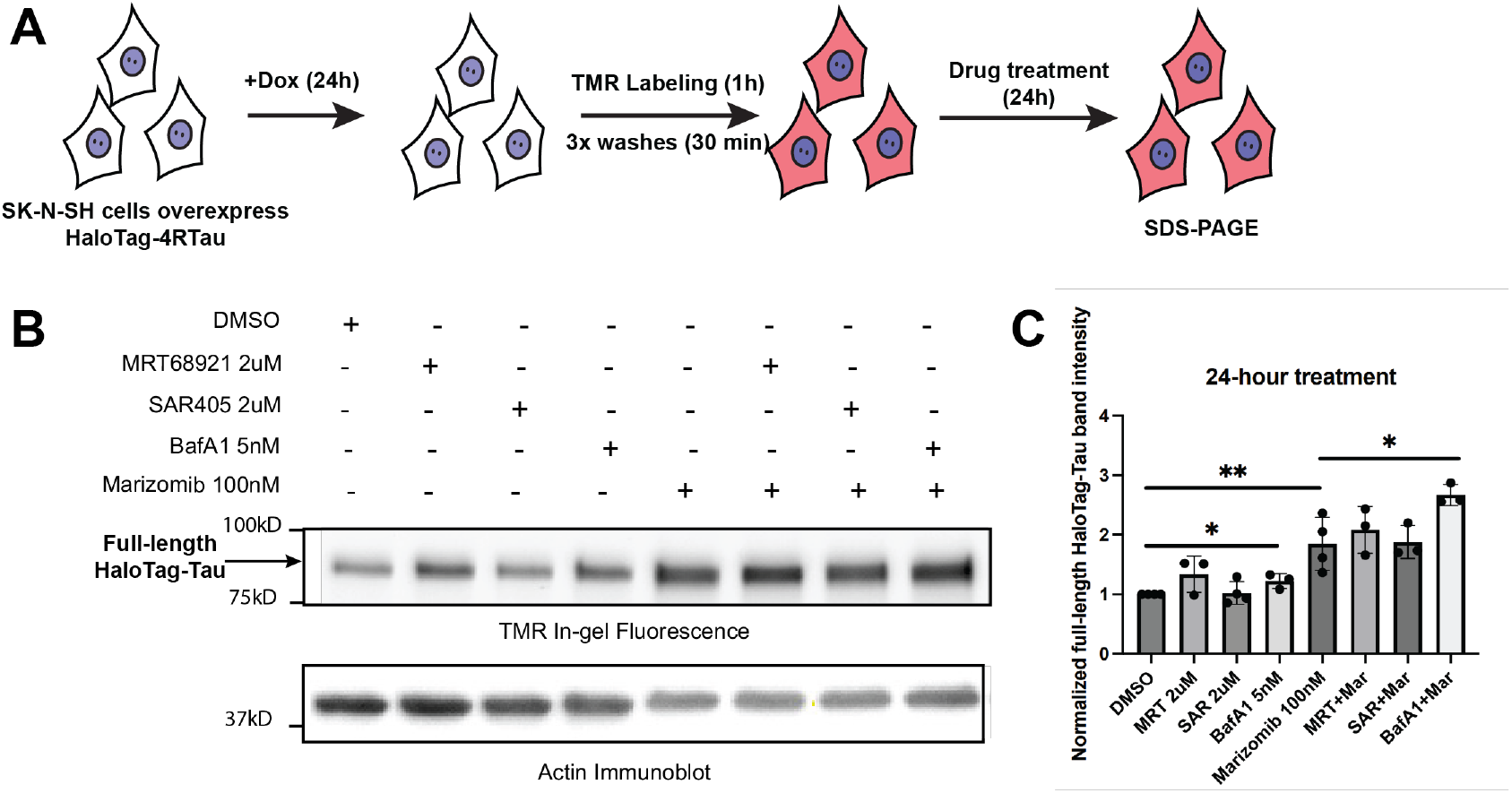
Determination of degradative pathways responsible for HaloTag–Tau turnover in neuroblastoma cells. (A) Schematic of the workflow for treating neuroblastoma cells overexpressing the HaloTag–4RTau sensor with drugs for SDS–PAGE analysis. (B) Representative SDS–PAGE in-gel fluorescence images TMR–HaloTag–4RTau under different treatment conditions. (C) Quantification and statistical analysis of band intensities of full-length TMR–HaloTag–Tau from in-gel fluorescence. Data are presented as mean ± SD. Statistical significance was determined using an unpaired Student’s t-test: *p < 0.05, **p < 0.01.

### Proteosome inhibition or lysosome deacidification accelerate HaloTag-Tau’s turnover via activation of caspases

We then tried the same pharmacological inhibition of proteasome and lysosome independently and in combination within iNeurons overexpressing the HaloTag-4RTau sensor. Surprisingly, all three treatments resulted in a significant decrease in full-length HaloTag–4R-Tau, as assessed by both TMR in-gel fluorescence and 4R-tau immunoblotting (Figure 4A–D). We measured caspase activity from iNeuron lysate after 24h-treatment with Marizomib and Bafilomycin A1 using the commercially available kit Caspase-Glo 3/7 Assay System, since caspases were reported to truncate Tau^20^ outside of proteasomal and lysosomal processing. It turned out that both proteasome and lysosome inhibition activated caspase activity (Figure 4E). To minimize caspase-mediated effects, we reduced the doses of marizomib and bafilomycin A1, but both treatments still induced caspase activation–mediated tau degradation (Figure 4F–J). We then shortened the treatment duration to 4 hours (instead of 24 hours) using higher doses (25 nM Marizomib and 100 nM Bafilomycin A1). Under these conditions, neither Marizomib nor Bafilomycin A1 had a significant effect on the intensity of the full-length TMR–HaloTag–4RTau band (Figure S3A–C). The combination of Marizomib and Bafilomycin A1 treatment slightly reduced the full-length TMR–HaloTag–4RTau band intensity, likely due to caspase activation. These results suggest that proteotoxicity from inhibiting protein degradative pathways contributes to tau cleavage by activating caspase activity.

**Figure 4.**
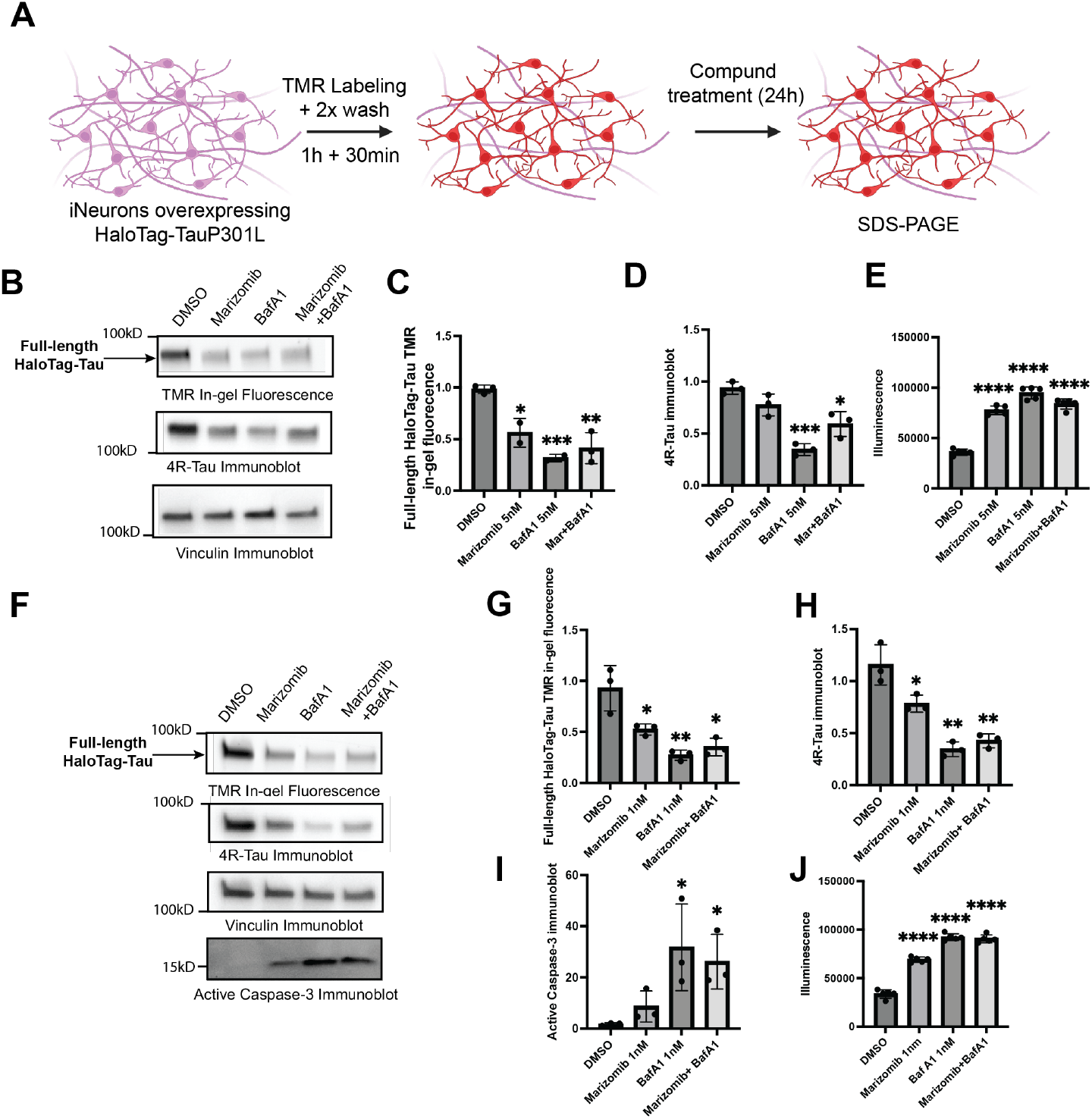
Evaluation of the degradative pathways and caspase activation in iNeurons expressing HaloTag–4RTau. (A) Workflow for treating iNeurons expressing the HaloTag–Tau sensor with proteasome and lysosome inhibitors for SDS–PAGE analysis. (B) Representative SDS–PAGE in-gel fluorescence images of TMR–HaloTag–Tau and immunoblots of 4R–Tau under different treatment conditions. (C–D) Quantification and statistical analysis of full-length TMR–HaloTag–Tau in-gel fluorescence and 4R–Tau immunoblot intensities normalized to Vinculin. Data are presented as mean ± SD; statistical significance was determined using an unpaired Student’s t-test: *p < 0.05, **p < 0.01, ***p < 0.001. (E) Quantification and statistical analysis of luminescence intensity from the Caspase-Glo 3/7 assay following drug treatment at indicated doses. Data are presented as mean ± SD; statistical significance was determined using an unpaired Student’s t-test: ****p < 0.0001. (F) Representative SDS–PAGE in-gel fluorescence images of TMR–HaloTag–Tau and immunoblots of 4R–Tau and active caspase-3 under different treatment conditions. (G–I) Quantification and statistical analysis of full-length TMR–HaloTag–Tau, 4R–Tau, and active caspase-3 immunoblot intensities normalized to Vinculin. Data are presented as mean ± SD; statistical significance was determined using an unpaired Student’s t-test: *p < 0.05, **p < 0.01. (J) Quantification and statistical analysis of luminescence intensity from the Caspase-Glo 3/7 assay following drug treatment at indicated doses. Data are presented as mean ± SD; statistical significance was determined using an unpaired Student’s t-test: ****p < 0.0001.

### Neddylation inhibition-mediated tau lowering effect is independent of caspase activation

Pevonedistat was reported to kill cancer cells by inducing apoptosis^21^, which can activate caspases, and caspase-mediated tau truncation has been implicated in tauopathy pathogenesis^20^. To determine whether Pevonedistat-mediated tau reduction involves caspase activation, we measured caspase activity from iNeuron lysate after 24h-treatment with Pevonedistat at various concentrations using the Caspase-Glo 3/7 Assay System. At concentrations (0.1uM and 1uM) efficient to reduce tau level (Figure 5A-C) for 24-hour treatment, Peveonedistat did not induce caspase activity (Figure 5D), with the proteasome inhibitor Marizomib and v-ATPase inhibitor Bafilomycin A1 serving as positive controls (Figure 5E). These results indicate that Pevonedistat-mediated tau clearance occurs independently of caspase activity.

**Figure 5.**
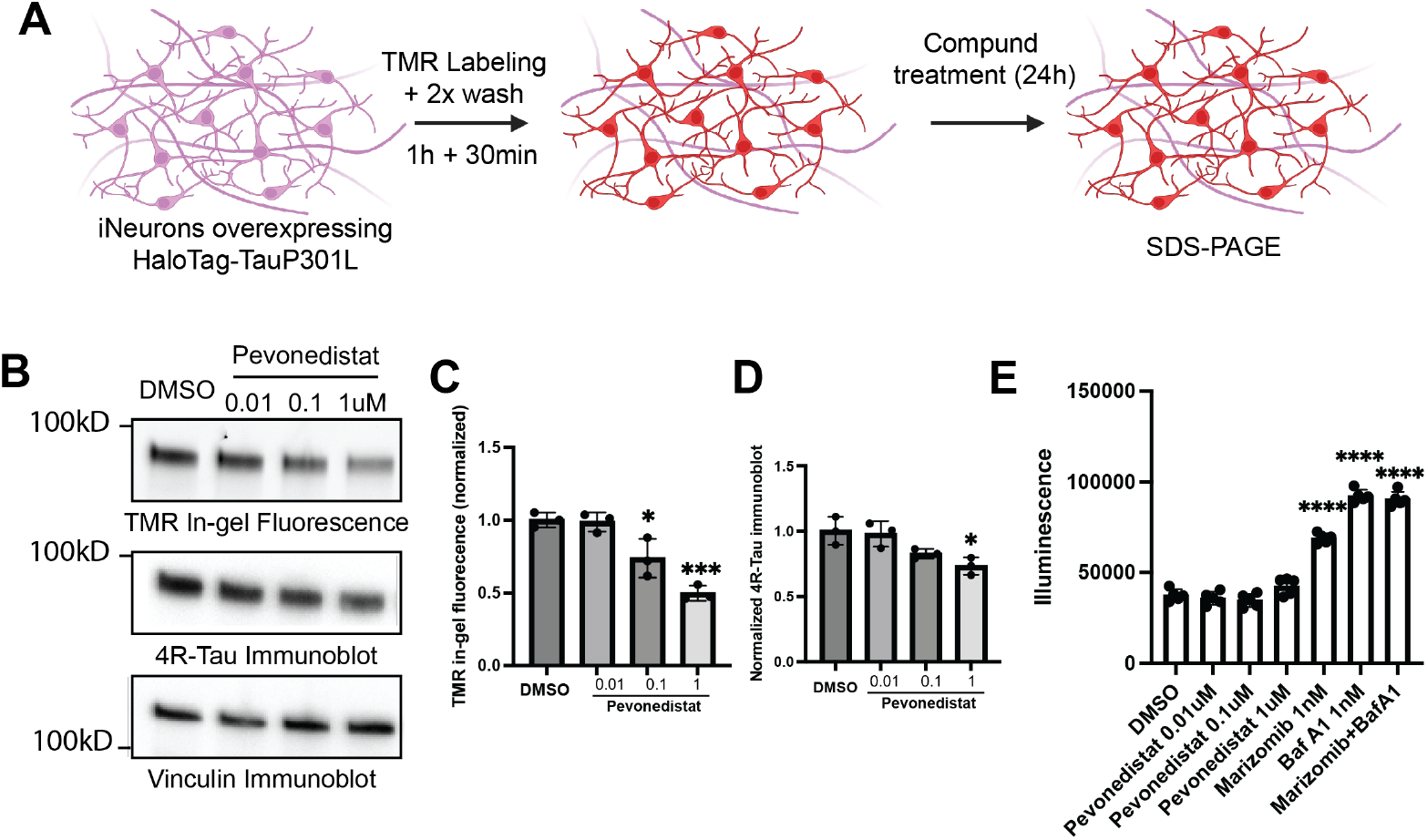
Reevaluation of dose dependence of Pevonedistat on HaloTag-Tau clearance and caspase activity in iNeurons. (A) Workflow for assessing Pevonedistat for HaloTag-Tau clearance and caspase activity in iNeurons expressing the HaloTag–Tau sensor after 24-hour drug treatment. (B) Representative full SDS–PAGE in-gel fluorescence images and immunoblots of TMR–HaloTag–Tau under different treatment conditions. (C-D) Quantification and statistical analysis of band intensities of full-length TMR– HaloTag–Tau from in-gel fluorescence and immunoblot. Data are presented as mean ± SD. Statistical significance was determined using an unpaired Student’s t-test: *p < 0.05, ***p < 0.001. (E) Quantification and statistical analysis of luminescence intensity from the Caspase-Glo 3/7 assay following treatment with different drugs at indicated doses. Data are presented as mean ± SD, and statistical significance was determined using an unpaired Student’s t-test: ****p < 0.0001.

### Neddylation inhibition-mediated tau lowering effect is likely through both the proteasome and autophagy-lysosome pathway

To further investigate the mechanism of neddylation inhibition–mediated tau reduction, iNeurons overexpressing HaloTag–Tau were treated with Pevonedistat (0.1 μM and 1 μM) for 24 hours, followed by unenriched proteomic analysis (Figure 6A). Compared with DMSO controls, BTB/Kelch proteins (BTBD2, BTBD10/KLHL9, KLHDC3) and F-box proteins (FBXO28, FBXO3) were significantly upregulated (Figure 6B–C). These proteins act as substrate adaptors for CRL3 and CRL1/SCF complexes, ^22, 23^ which are substrates of Neddylation. Because neddylation inhibition inactivates CRLs^24^, their upregulation likely reflects a compensatory response to CRL inactivation instead of insufficient substrates degradation as we did not observe accumulated CRLs substrates. We also observed upregulation of AMBRA1, a CRL substrate adaptor that promotes autophagy by activating the Beclin1–Vps34 PI3K complex^25, 26^. This suggests that Pevonedistat may facilitate lysosomal degradation of HaloTag–Tau via AMBRA1-dependent autophagy, potentially complementing previously reported mTOR-dependent pathways^18^. Notably, COP1, a RING-type E3 ligase unrelated to CRLs, was also upregulated. This indicates that Pevonedistat may additionally enhance compensatory proteasome activity, contributing to increased turnover of soluble tau. Together, these findings support a model in which neddylation inhibition triggers adaptive changes in both proteasome and autophagy pathways to promote tau clearance.

**Figure 6.**
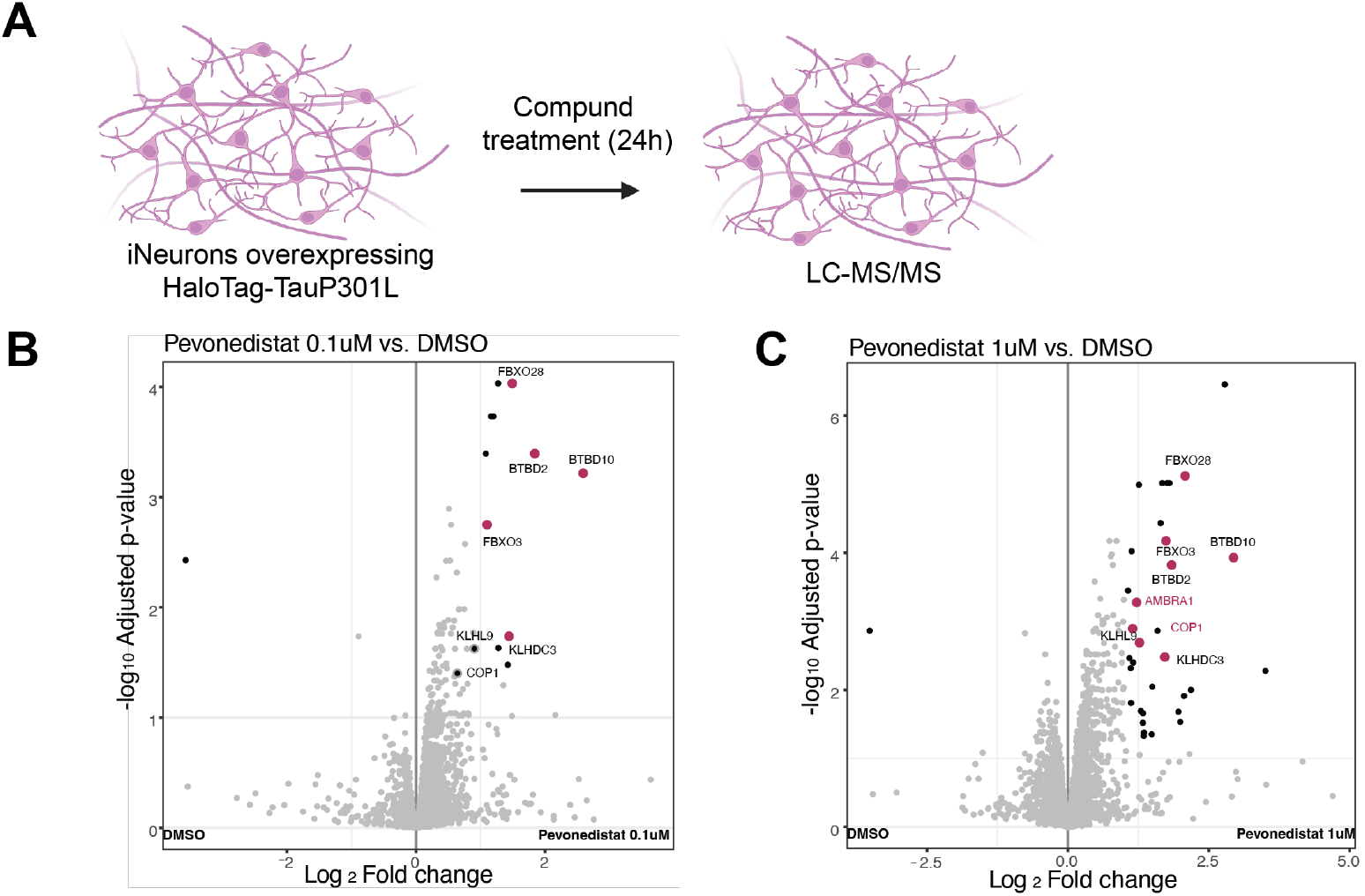
Identification of major degradative pathways for HaloTag–Tau turnover in iNeurons and proteomic changes induced by Pevonedistat. (A) Schematic of the workflow for treating iNeurons expressing HaloTag–Tau sensor with Pevonedistat for 24 hours prior to mass spectrometry-based proteomic analysis. (B–C) Volcano plots showing proteins with log2 fold changes (log2 FC) and adjusted p-values following Pevonedistat treatment (0.1 μM and 1 μM) versus DMSO control. Cutoffs for significance were set at |log2 FC| ≥ 1 and adjusted p-value ≤ 0.05.

## Discussion

In this study, we developed a tau sensor that enables sensitive detection of lysosomal localization and overall cellular turnover of tau via pulse-chase experiments. HaloTag-tagged Tau retains its ability to bind microtubules and exhibits a half-life similar to that of endogenous Tau reported previously.^11^ Using this sensor, we screened a library of proteostasis modulators to investigate how perturbation of proteostasis network function affects HaloTag-Tau and to identify druggable pathways that enhance its degradation. Among the positive hits promoting lysosomal localization, we confirmed that neddylation inhibition accelerates the degradation of soluble HaloTag–Tau. Mechanistic studies further indicate that the both autophagy-lysosome and proteasome pathways are responsible for soluble tau clearance, and that neddylation inhibition–mediated tau degradation likely occurs via compensatory proteasomal activity and autophagy activation, as evidenced by both the lysosomal translocalization effect and proteomic analysis.

The innovative aspect of this study lies in the development of a tau sensor capable of detecting lysosomal localization of tau with high sensitivity using a pulse–chase approach. Another novel feature is the screening of a small-molecule library targeting proteostasis network function, as protein aggregates such as tau often arise from disrupted proteostasis. Our results show that inhibition of several pathways within the proteostasis network can either block lysosomal localization of tau or slow the cellular turnover of HaloTag–Tau. The tau-lowering effect of neddylation inhibition and the tau-accumulating effect of mTOR inhibition are consistent with the effects observed in previous studies using genetic knockdown of these targets^27^.

One limitation of this study is that the mechanism underlying pevonedistat-mediated tau clearance was not fully explored—for example, through genetic knockdown of key components of the neddylation pathway or co-treatment with a proteasome inhibitor. Previous studies have reported that genetic knockdown of neddylation reduces soluble tau levels, and co-treatment of pevonedistat with proteasome inhibition further decreases soluble tau, likely via caspase activation (or activation of the autophagy lysosome pathway by independent mechanisms, as it has been reported that proteasome inhibition activates autophagy)^8^. Our data indicate that proteotoxicity from proteasome inhibition or lysosome deacidification can activate caspases, cleaving full-length tau. These findings underscore the indirect effects that proteasome and lysosome dysfunction can have on tau pathology and highlight the need for careful interpretation of tau turnover assays under conditions that perturb these pathways. Further studies are needed to fully understand the portioning of TMR-HaloTag–Tau through the proteasome and the lysosome upon neddylation inhibition or with other pharmacological treatments.

## Acknowledgements

This work was supported by the National Institute on Aging (RF1AG073418) and the Rainwater Charitable Foundation. We greatly appreciate Emily P. Bentley’s expert editorial feedback.

## Methods and materials

### CRISPR knock in of NGN2 in iPSCs

Following the protocol published by the Michael Ward Lab, sgRNA targeting the CLYBL safe-harbor locus (sequence: ATGTTGGAAGGATGAGGAAA), together with Cas9 RNP (#309602800 from IDT) and donor DNA (CLYBL-TO-hNGN2-BSD-mApple, a gift from Michael Ward, Addgene plasmid #124229;http://n2t.net/addgene:124229;RRID:Addgene_124229), was delivered into human iPSCs (Aurora BioLabs). Single clones were isolated by fluorescence-activated cell sorting (FACS) based on mApple fluorescence and subsequently karyotyped by WiCell Characterization Services. Subclones with the correct chromosome number and normal structure (Figure S1F) were used for all subsequent experiments.

### Stable HaloTag/HaloTag-TauP301L sensor iPSC line

HaloTag and HaloTag–TauP301L sequences were cloned into a third-generation TetON-inducible lentiviral vector and packaged into lentivirus (VectorBuilder). NGN2-engineered iPSCs were transduced with the lentivirus at an MOI of 2. Sensor-positive cells were enriched by antibiotic selection (puromycin resistance) and validated by visualizing HaloTag–Tau in iNeurons using the fluorescent HaloTag ligand TMR (#G8281, Promega) or the fluorogenic HaloTag ligand JF646 (#GA1120, Promega), as described in detail in the next section. Overexpressed HaloTag and HaloTag–TauP301L proteins were further confirmed by SDS–PAGE and Western blotting (Figure 2D, 2H).

### iNeuron differentiation

Sensor-positive cells were detached and individualized using accutase (#07920 from Stemcell Technologies Inc.) and reseeded onto Matrix Gel-coated plates (#CB40234, Corning) in mTeSR+ media (#100-0276, Stemcell Technologies Inc.) supplemented with 10 μM ROCK inhibitor (Y-27632 dihydrochloride, #HY-10071, MedChemExpress). The ROCK inhibitor was removed on the second day, and doxycycline (2 μg/mL, #HY-N0565, MedChemExpress) was added to the media, which was maintained for 2 days. Cells were then lifted using accutase and seeded onto poly-Ornithine-coated plates (PhenoPlate 384-well, #6057320, Revvity, or chamber slides, #81817, iBidi) in neuron maturation media (NMM), consisting of a 1:1 mixture of NeuroBasal media (#10888022, Gibco) and DMEM/F12 (#21041025, Gibco), supplemented with N-2 (#17502048, Gibco), B-27 (#17504044, Gibco), laminin (0.5 μg/mL, #23017015, Gibco), and doxycycline (2 μg/mL). NMM was refreshed every other day for 2 weeks.

### Pulse-chase experiment for live/fix-cell imaging and SDS-PAGE

Mature iNeurons overexpressing HaloTag or HaloTag–Tau were labeled with 1 μM HaloTag ligand tetramethylrhodamine (TMR, #G8281, Promega) or 50 nM JF646 (#GA1120, Promega) in neuron maturation media (NMM, without doxycycline) for 1 hour. Cells were then washed twice with NMM (without doxycycline, containing 5 μM HaloTag blocker, non-fluorescent HaloTag ligand 1-chloro-6-(2-propoxyethoxy)hexane, #F83341-25MG, AstaTech, Inc.) and incubated for 30 minutes before applying fresh NMM (without doxycycline, containing compounds and 5 μM HaloTag blocker) for the desired treatment duration.

For live-cell imaging, cells were directly placed in a Zeiss Celldiscoverer 7. For fixed-cell imaging, cells were fixed with 4% paraformaldehyde and immunostained with MAP2 (#ab183830, Abcam), NeuN (#ab104225, Abcam), or Tuj-1 (#ab18207, Abcam).

For SDS–PAGE analysis, cells were lysed in RIPA buffer supplemented with protease and phosphatase inhibitors (#PI87785, Thermo Fisher Scientific) for 30 minutes on ice. Protein concentrations were normalized using a BCA assay (#PI23250, Thermo Fisher Scientific) and samples were boiled in SDS sample buffer for 5 minutes. In-gel fluorescence was detected using a ChemiDoc imager (Bio-Rad). HaloTag antibody (#G9211 prom Promaga), 4R-Tau antibody (#30328 from Cell Signaling Technology), Actin antibody (#4967 from Cell Signaling Technology), and Vinculin antibody (#4650 from Cell Signaling Technology), were used for immunoblotting.

### Image analysis and quantification

Acquired images were analyzed using Arivis (v4.20, Zeiss). Puncta and neuronal soma segmentation were performed with the machine-learning module of the analysis pipeline, using training datasets generated by manually annotating areas of interest at random. The analysis pipeline was applied in batch mode to generate quantification data for each well. The resulting data files were used to generate heatmaps using Python code, which has been deposited on GitHub (DOI: 10.5281/zenodo.17903202).

### Mass spectrometry sample prep

Mature iNeurons overexpressing HaloTag–Tau and treated with pevonedistat for 24 hours were detached by gentle shaking and washed twice with PBS. Cell pellets were lysed in 8 M urea buffer containing 75 mM NaCl, 50 mM HEPES (pH 7.5), protease inhibitors (cOmplete™, Mini, EDTA-free, #11836170001, Sigma), and phosphatase inhibitors (PhosSTOP, #04906837001, Sigma) for 15 minutes on ice. Lysates were collected after centrifugation at 13,000 rpm for 15 minutes at 4°C.

BCA-normalized lysates were treated with 5 mM TCEP for 30 minutes at room temperature for disulfide reduction, followed by 14 mM iodoacetamide for 30 minutes at room temperature in the dark for cysteine alkylation. Proteins were precipitated using methanol/chloroform/water (4:1:3). Precipitated proteins were digested with Trypsin–LysC (Thermo Scientific) at a 1:100 enzyme-to-protein ratio in 200 mM EPPS buffer (pH 8.5) overnight at 37°C in a thermomixer shaking at 1,500 rpm.

The digested peptide mixture was acidified with 0.1% TFA to pH <2 and desalted using Pierce Peptide Desalting Spin Columns. The eluted peptides were dried in a speedvac overnight and resuspended in 0.1% formic acid for concentration measurement by absorbance at 205 nm. One microgram of peptides was injected into an Orbitrap Exploris 480 mass spectrometer, and data were acquired using data-independent acquisition (DIA).

### Mass spectrometry data analysis

Raw mass spectrometry data were processed using FragPipe (v22.0) with MSFragger for database searching, Philosopher for false discovery rate (FDR) filtering, and IonQuant for quantification. Spectra were searched against the UniProt human proteome, including common contaminants and decoy sequences. MSFragger searches assumed trypsin digestion with up to two missed cleavages, ±20 ppm precursor tolerance, 0.02 Da fragment tolerance, fixed carbamidomethylation of cysteines, and variable methionine oxidation and N-terminal acetylation. IonQuant performed label-free quantification with match-between-runs enabled and default normalization. Protein intensities were derived from unique peptides and quantified by DIA-NN.

Quantified protein data were exported to R, log2-transformed, and filtered to remove missing values. Differential abundance analysis was performed using linear modeling with Benjamini– Hochberg correction for multiple testing. Results were visualized using volcano plots.

**Figure S1.**
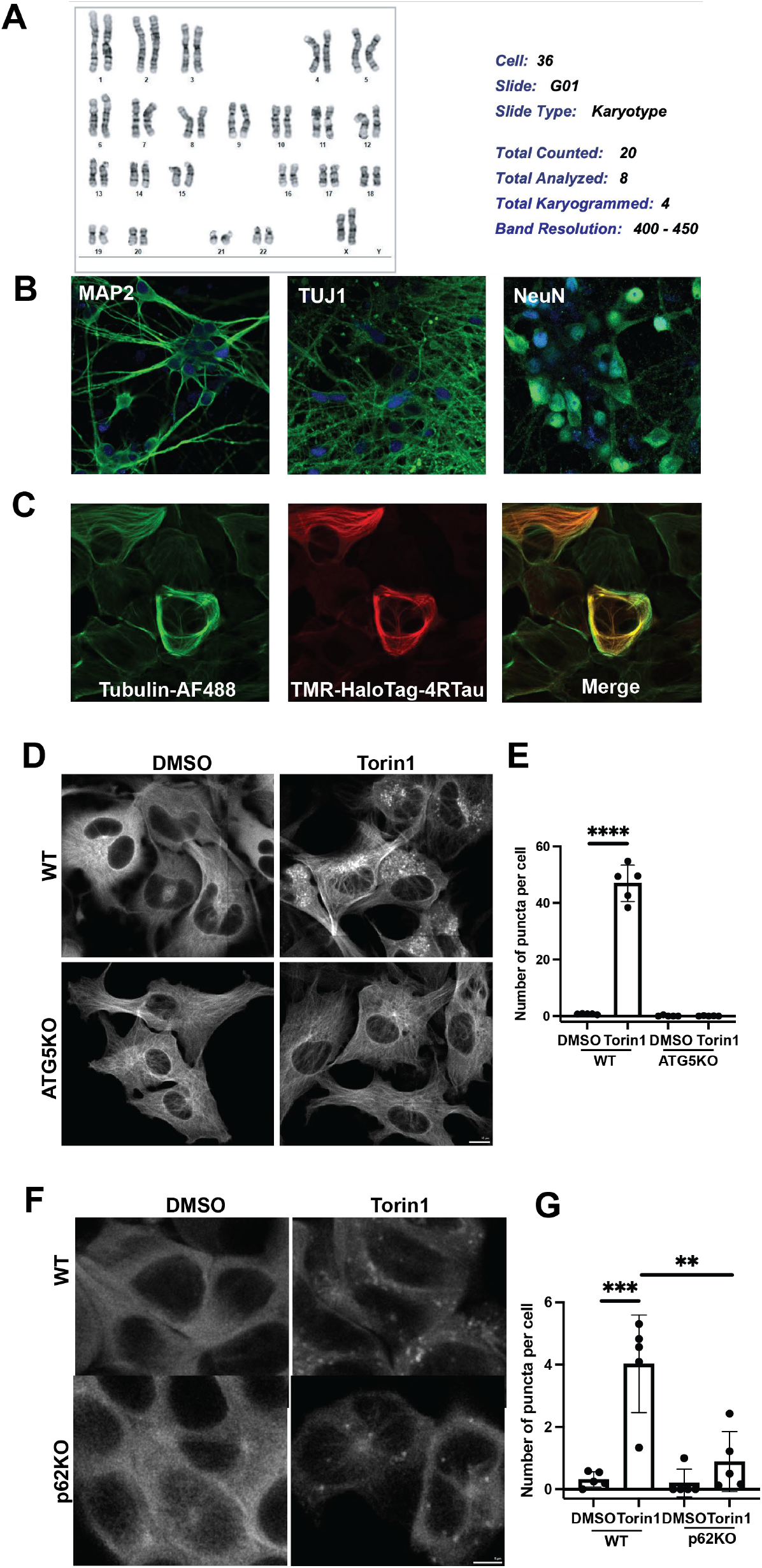
Characterization of iNeurons and autophagy-dependence of HaloTag–Tau lysosomal localization. (A) Immunostaining of neuronal markers MAP2, TUJ1, and NeuN in iNeurons after 2 weeks of differentiation. (B) Chromosome analysis of iPSCs engineered with NGN2 and HaloTag–Tau, showing normal karyotype. (C) Representative images of the colocalization of TMR-HaloTag-4RTau with microtubules, immunostained by Tubulin-AF488 antibody. (D)Representative images of U251 cells transduced with HaloTag–Tau with or without ATG5 knockout (ATG5KO), treated with DMSO or Torin1 (500 nM) for 24 hours. Scale bar, 10 μm. (E) Quantification and statistical analysis of TMR–HaloTag–Tau puncta per cell in U251 cells. Data are presented as mean ± SD; statistical significance was determined using an unpaired Student’s t-test: ****p < 0.0001. (F) Representative images of HEK293 cells transduced with HaloTag–Tau with or without p62 knockout (p62KO), treated with DMSO or Torin1 (500 nM) for 24 hours. Scale bar, 5 μm. (G) Quantification and statistical analysis of TMR–HaloTag–Tau puncta per cell in HEK293 cells. Data are presented as mean ± SD; statistical significance was determined using an unpaired Student’s t-test: **p < 0.01, ***p < 0.001.

**Figure S2.**
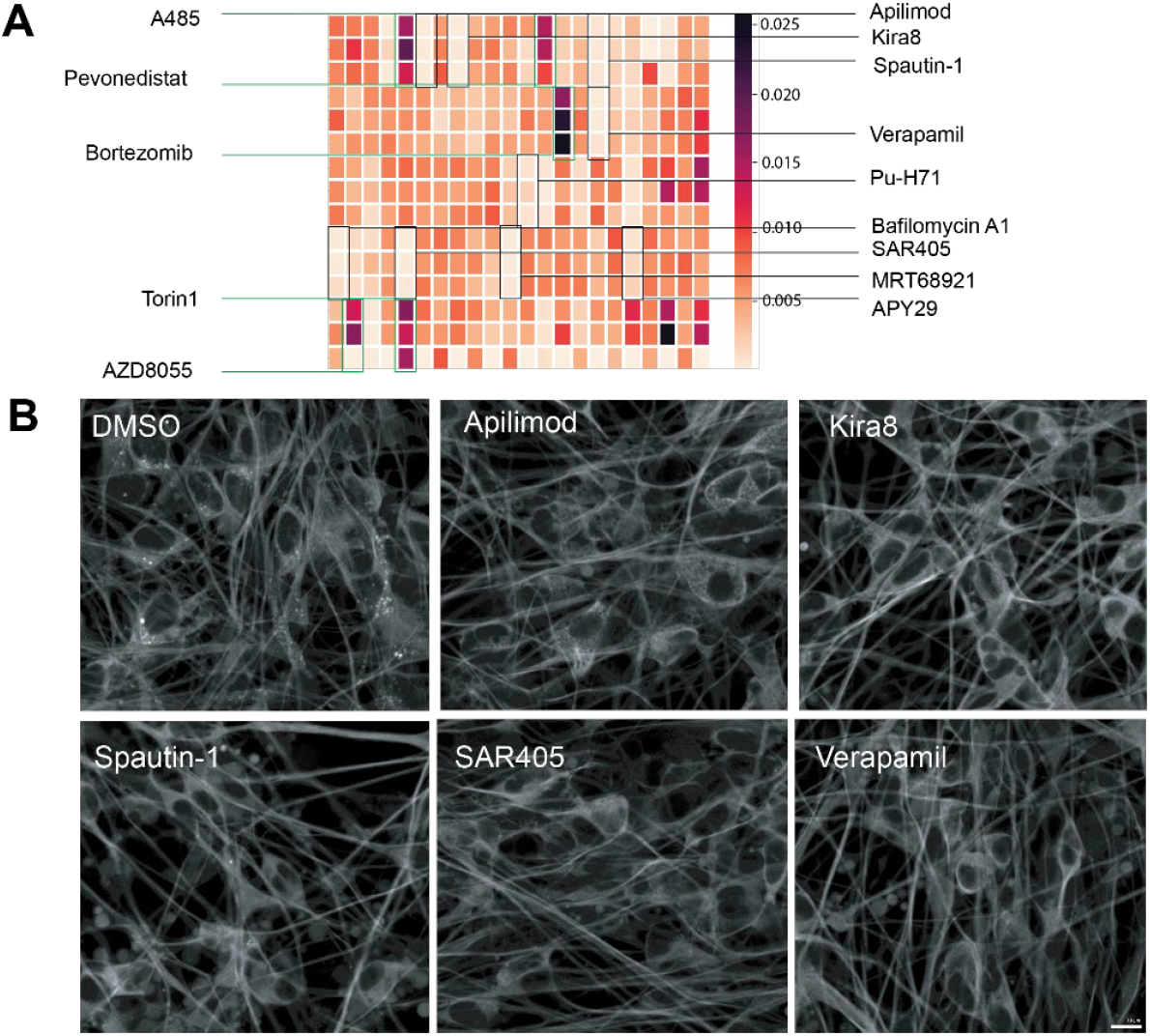
High-content screening of proteostasis modulators using HaloTag–Tau sensor in iNeurons. (A) Heatmap showing the number of puncta per neuronal area as measured by the HaloTag– Tau sensor in iNeurons treated with proteostasis modulators (triplicate wells, 10 μM) for 24 hours. Darker colors indicate a higher number of puncta per area. (B) Representative images of negative hits that inhibit lysosomal translocation of HaloTag–Tau. Scale bar, 10 μm.

**Figure S3.**
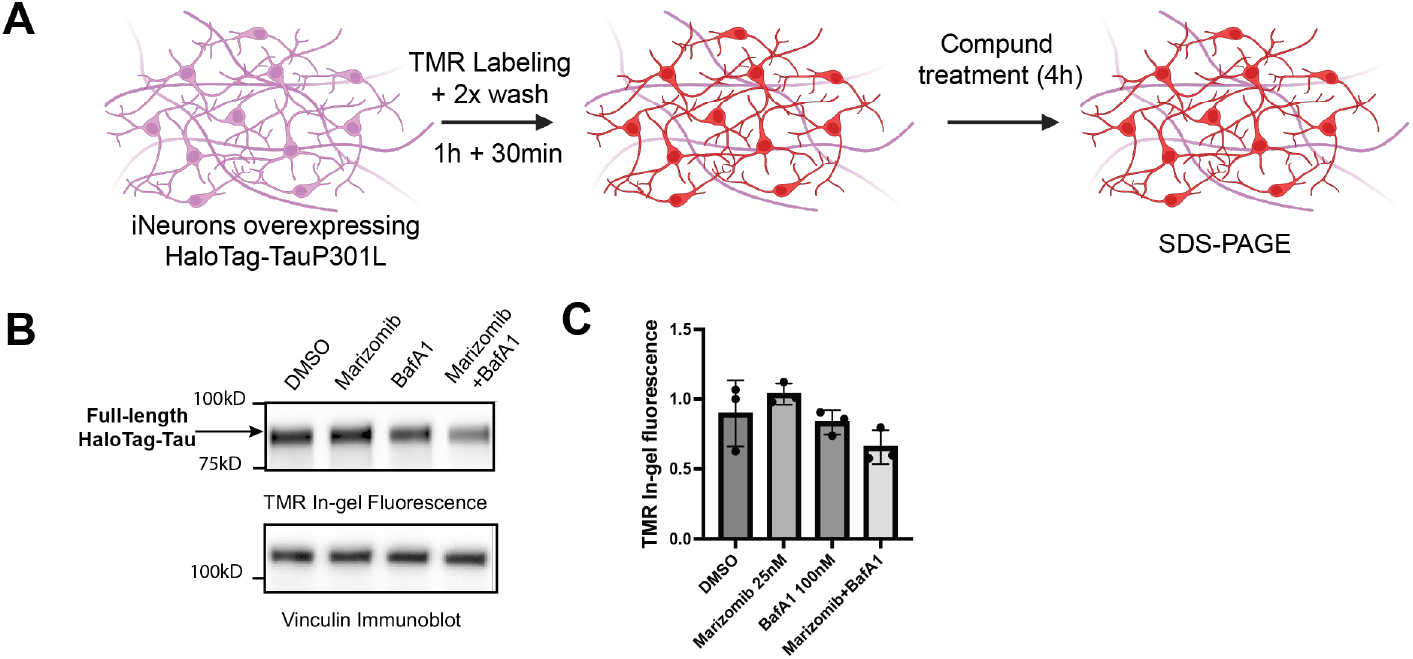
Evaluation of the degradative pathways in iNeurons expressing HaloTag–4RTau. (A) Schematic of the workflow for treating iNeurons expressing the HaloTag–Tau sensor with proteasome and lysosome inhibitors for SDS–PAGE analysis. (B) Representative full SDS–PAGE in-gel fluorescence images of TMR–HaloTag–Tau under different treatment conditions. (C) Quantification and statistical analysis of full-length TMR–HaloTag–Tau band intensities from in-gel fluorescence. Data are presented as mean ± SD, and statistical significance was determined using an unpaired Student’s t-test.

**Figure S4.**
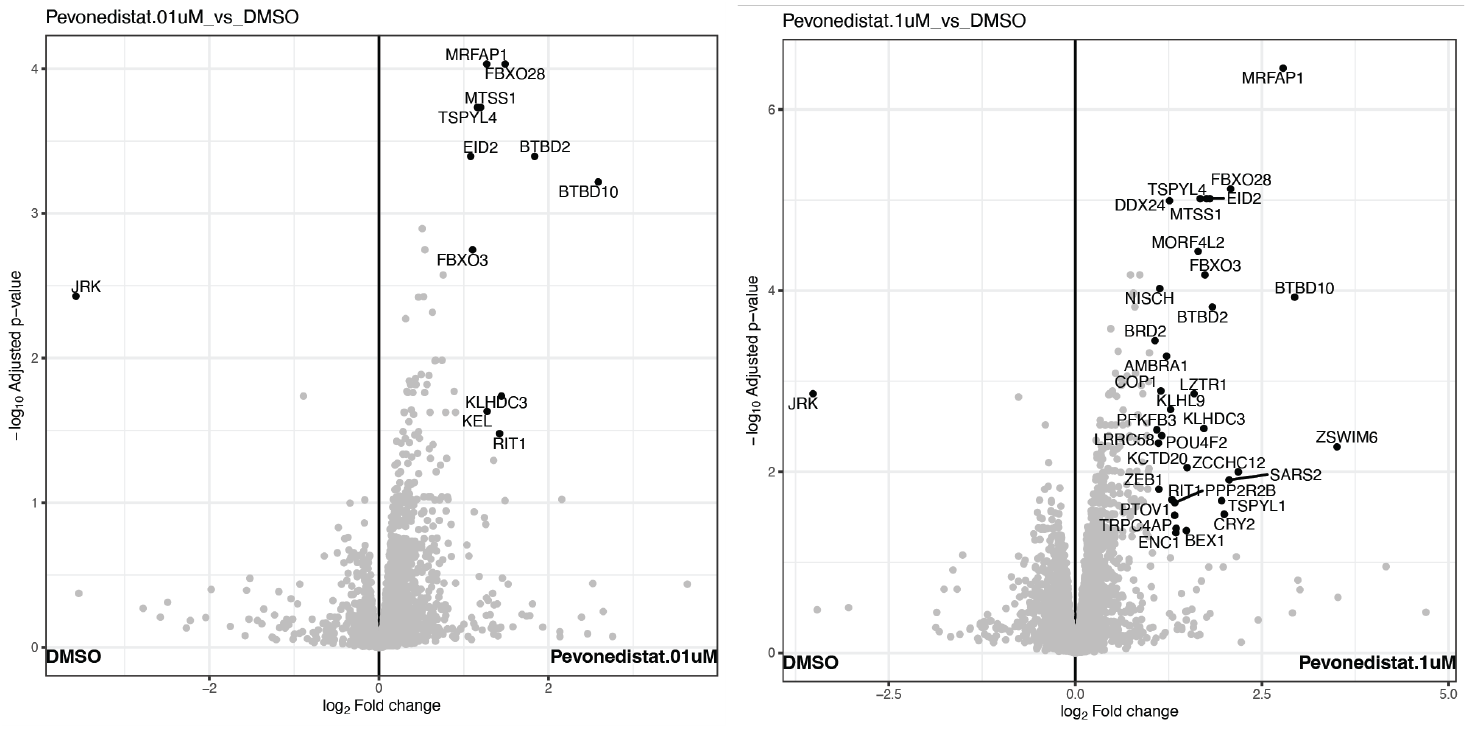
Proteomic changes induced by Pevonedistat in iNeurons. Volcano plots showing proteins with log2 fold changes (log2 FC) and adjusted p-values following Pevonedistat treatment (0.1 μM and 1 μM) compared with DMSO control. Cutoffs for significance were set at |log2 FC| ≥ 1 and adjusted p-value ≤ 0.05.

